# A real-time compression library for microscopy images

**DOI:** 10.1101/164624

**Authors:** Bálint Balázs, Joran Deschamps, Marvin Albert, Jonas Ries, Lars Hufnagel

**Affiliations:** Cell Biology and Biophysics Unit, EMBL Heidelberg, Germany; Faculty of Information Technology and Bionics, Pázmány Péter Catholic University, Budapest, Hungary; Developmental Biology Unit, EMBL Heidelberg, Germany.

## Abstract

Fluorescence imaging techniques such as single molecule localization microscopy, high-content screening and light-sheet microscopy are producing ever-larger datasets, which poses increasing challenges in data handling and data sharing. Here, we introduce a real-time compression library that allows for very fast (beyond 1 GB/s) compression and de-compression of microscopy datasets during acquisition. In addition to an efficient lossless mode, our algorithm also includes a lossy option, which limits pixel deviations to the intrinsic noise level of the image and yields compression ratio of up to 100-fold. We present a detailed performance analysis of the different compression modes for various biological samples and imaging modalities.

## Main

Advancements in fluorescence microscopy technologies such as in high-content screening [1]–[3], light-sheet microscopy [4]–[8], and single molecule localization microscopy [9]–[11] opened new perspectives in biology by increasing the speed of imaging, the number of specimens or the resolution of the observed structures. Even though these methods bring undeniable advantages, the data production speed and experiment sizes (**
Fig. 1a, Supplementary Table 1**) are increasing in such a fast pace that in many cases data handling quickly becomes a bottleneck for new discoveries [12]–[14]. A straightforward solution to this problem is to perform image compression. Nonetheless, this typically implies incompatibilities with certain software packages, slow compression speed, and only moderate file size reduction for lossless methods. Although the compression ratio (original size / compressed size) can be substantially increased with lossy compression algorithms, their use is often discouraged [15] as the degree of information loss heavily depends on the image content and cannot be explicitly controlled.

**Figure 1.**
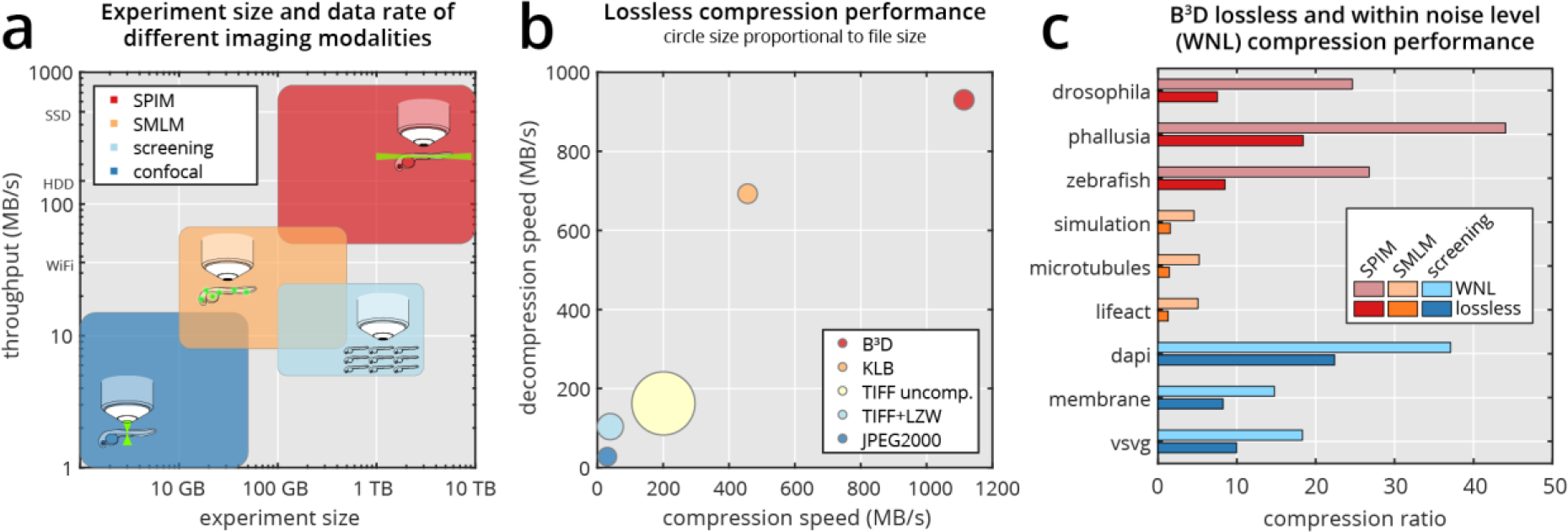
Comparing image compression methods for high-speed microscopy. (a) Comparison of single-plane illumination microscopy (SPIM, red rectangle), high-content screening (light blue), single molecule localization microscopy (SMLM, orange) and confocal microscopy (blue) by typical experiment size and data production rate (see also Supplementary Table 1). (b) Performance comparison of our B³D compression algorithm (red circle) vs. KLB (orange), uncompressed TIFF (light yellow), LZW compressed TIFF (light blue) and JPEG2000 (blue) regarding write speed (horizontal axis), read speed (vertical axis) and file size (circle size). (c) Lossless and WNL compression ratios for SPIM, SMLM and screening microscopy datasets. (Supplementary Table 3).

To address these challenges, we developed a new compression library called B^3^D, which is capable of extremely fast compression and decompression of large microscopy datasets. Our library is built on the CUDA architecture [16] for GPU-based compression, which not only enables high processing speed, but also relieves load on the central processing unit, allowing compression directly during image acquisition. The algorithm has two main components. First, a prediction is made for each pixel based on the neighboring pixel values, and second, the prediction errors are run-length and Huffman encoded to effectively reduce the data size (**Supplementary Note** and **Supplementary Fig. 1**). We compared our algorithm’s performance with TIFF (LZW), JPEG2000, and the speed-optimized KLB [17] by measuring compression speed, decompression speed and resulting file size (**
Fig. 1b
**). Only B^³^D is capable of handling the sustained high data rate of modern sCMOS cameras typically used in light-sheet microscopy, while still maintaining compression ratios comparable to more complex, but much slower algorithms (**Supplementary Table 2**).

Lossless compression is fundamentally limited by the algorithmic entropy of the data, and further reduction in size is only possible through loss of information. Most lossy compression algorithms (such as JPEG) compress images by preserving only those structures recognized by the human visual system [18], making compression artifacts imperceptible to the eye. Scientific images, however, represent sets of quantitative measurements and therefore their compression should instead preserve the numerical values of all pixel intensities within their uncertainties. Pixel values are subject to random noise that mainly consist of the photon shot noise and the camera read noise. Commonly used lossy compression algorithms often perform a quantization step in Fourier or wavelet space. Therefore, the deviation introduced by the compression at the single pixel level cannot be controlled. We developed a noise dependent lossy compression algorithm which processes each pixel individually and exploits the natural variability of pixel values. As a result, the user can specify the maximally tolerated pixel error in proportions of the inherent noise. Extending our lossless compression scheme, this is achieved by adding a variance stabilization step before the prediction, and quantizing the prediction residuals before the Huffman coding (**Supplementary Note**). Alternatively, the quantization and prediction steps can be swapped, which can be more suitable for methods that are sensitive to small scale pixel correlations, such as localization microscopy, albeit at the expense of compression ratio (**Supplementary Fig. 2**). We define the mode where the quantization step is equal to the noise (*q*=1*σ*) as within noise level (WNL) compression. Using this method, the compression ratio massively increases for all imaging modalities compared to the lossless mode (**
Fig. 1c
**) without any apparent loss in image quality (**Supplementary Fig. 3**). Furthermore, the average compression error is considerably smaller than the image noise itself (**Supplementary Fig. 4**).

To see how this noise-dependent compression affects common imaging pipelines, we tested the effect of different levels of compression on 3D nucleus and membrane segmentation in light-sheet microscopy, and on single-molecule localization accuracy in superresolution microscopy. First, we imaged a *Drosophila melanogaster* embryo expressing an H2Av-mCherry nuclear marker in a MuVi-SPIM setup [6] and segmented the nuclei (**
Fig. 2a
** and **Online Methods**). Then we performed noise dependent compression at various quality levels and calculated the segmentation overlap compared to the uncompressed stack (**Online Methods**). At WNL compression (*q*=1*σ*) the segmentation overlap is almost perfect (**
Fig. 2b
**) with an overlap score of 0.996. Even when increasing the quantization step to 4*σ* (**
Fig. 2c
**) the overlap score stays at 0.98 and only drops below 0.97 when the compression ratio is already above 120 (quantization step of 5σ, **
Fig. 2d
**). We got similar results for a membrane segmentation pipeline that is used with *Phallusia mammillata* embryos (**Online Methods** and **Supplementary Fig. 5**).

**Figure 2.**
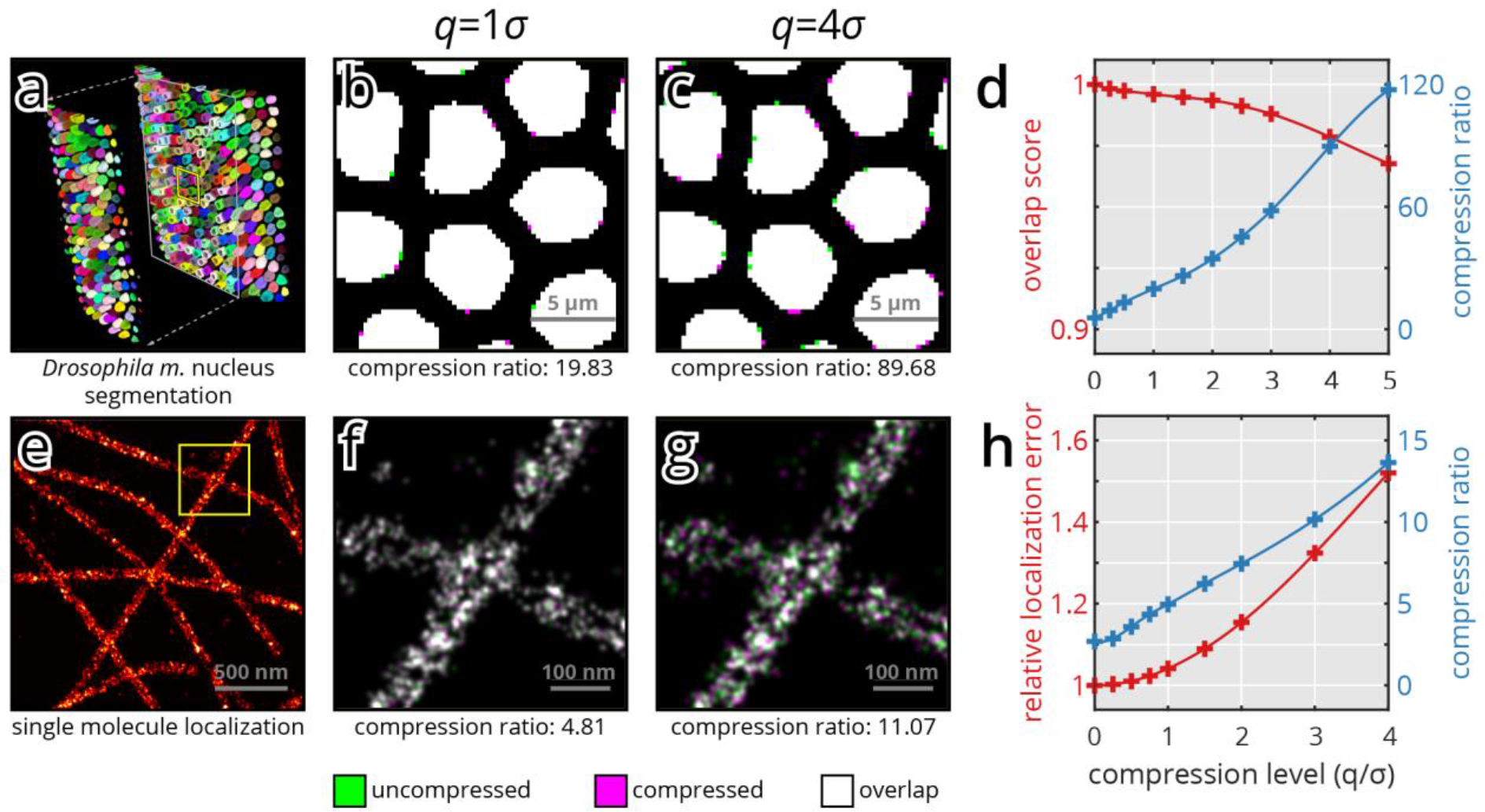
Effect of noise dependent lossy compression on image analysis outcome. (**a–h**) Influence of noise dependent lossy compression on 3D nucleus segmentation. A *Drosophila m*. embryo expressing H2Av-mCherry nuclear marker was imaged in MuVi-SPIM [7], and 3D nucleus segmentation was performed (Online Methods) (**a**). The raw data was subsequently compressed at increasingly higher compression levels, and segmented based on the training of the uncompressed data. To visualize segmentation mismatch, the results of the uncompressed (green) and compressed (magenta) datasets are overlaid in a single image (**b**, c; overlap in white). Representative compression levels were chosen at two different multiples of the photon shot noise, at *q*=1*σ* (**b**) and *q*=4*σ* (**c**). For all compression levels the segmentation overlap score (Online Methods) was calculated and is plotted in (**g**) along with the achieved compression ratios. (**e–h**) Influence of noise dependent lossy compression on single-molecule localization. Microtubules, immunolabeled with Alexa Fluor 647 were imaged by SMLM (**h**). The raw data was compressed at increasingly higher compression levels, and localized using the same settings as the uncompressed data. To visualize localization mismatch, the results of the uncompressed (green) and compressed (magenta) datasets are overlaid in a single image (**f**, **g**; overlap in white). Two representative compression levels were chosen at *q*=1*σ* (**f**) and *q*=4*σ* (**g**). To assess the effects of compression on localization precision, a simulated dataset with known emitter positions was compressed at various levels. For all compression levels the relative localization error (normalized to the Cramér–Rao lower bound) was calculated and is plotted in (**h**) along with the achieved compression factors.

Next, we evaluated our compression algorithm in the context of single molecule localization, and measured how the localization precision is affected by an increasing compression ratio. We compressed a single-molecule localization microscopy (SMLM) dataset of immuno-detected microtubules (**
Fig. 2e
**) with increasing compression levels. For WNL compression (*q*=1*σ*) no deterioration of the image was visible (**
Fig. 2f
**), and even for the case of *q*=4*σ* the compression induced errors were much smaller than the resolvable features (**
Fig. 2g
**). To quantify the impact of compression on the localization error, we used a simulated dataset (**Online Methods**) and compared the localization output of different compression levels to the ground truth (**
Fig. 2h
**). Lossless compression resulted in a compression ratio of 2.7, whereas WNL compression reached a compression ratio of 5.0, while increasing the localization error by only 4%. This also coincides with the theoretical increase of image noise (**Supplementary Note**). Furthermore, the increase in localization error was not dependent on the signal to background noise ratio (**Supplementary Fig. 6**).

Our algorithm is implemented in C++, and allows for easy integration through an API with various programming languages. The library was tested on Linux (Ubuntu 16.04) and Windows (10). Additionally, we implemented a filter plugin for HDF5 which enables a seamless integration in all software packages that are supporting the native HDF5 library, such as Matlab, Python, Imaris, or Ilastik. Because of its versatility, HDF5 has emerged as the *de facto* standard in the open source light sheet microscopy field, and is also the basis for the widely used BigDataViewer [19] in Fiji [20]. When loading a B^3^D compressed image in an HDF5 enabled application, the library automatically calls our filter plugin, decompresses the image on the GPU, and copies it back into CPU memory. Due to B^3^D’s efficient compression and its high decompression speed, loading data is often accelerated: For a state-of-the-art hard drive with 200 MB/s bandwidth, loading a 2 GB uncompressed 3D stack of images takes about 10 seconds. With an average compression ratio of 20 fold in the WNL mode, the loading time is reduced to 0.5 seconds followed by 2 seconds of decompression, which yields a factor of four speed-up. It is also worth to note, that the achieved WNL compression reduces the camera data rate to below 40 MB/s, well below the 1 Gb/s Ethernet standard. This enables to use current network infrastructure to move data to long term storage and even makes the use of cloud services possible. Altogether, B^3^D, our efficient GPU-based image compression library allows for exceptionally fast compression speed and greatly increases compression ratio with its WNL scheme, offering a versatile tool that can be easily tailored to any high-speed microscopy environment.

## Online Methods

### Compression benchmarking

For all presented benchmarks, TIFF and JPEG2000 performance was measured through MATLAB’s *imwrite* and *imread* functions, while KLB and B^³^D performance was measured in C++. All benchmarks were run on a computer featuring 32 processing cores (2×Intel Xeon E5-2620), 128 GB RAM and an NVIDIA GeForce GTX 970 graphics processing unit. Read and write measurements were performed in RAM to minimize I/O overhead, and are an average of 5 runs.

### Light-sheet imaging

*Drosophila melanogaster* embryos were imaged in our MuVi-SPIM setup [6] using the electronic confocal slit detection (eCSD) [21]. Embryos were collected on an agar juice plate, and dechorionated in 50% bleach solution for 1 min. The embryos were then mounted in a shortened glass capillary (Brand 100 μl) filled with 0.8% GelRite (Sigma-Aldrich), and pushed out of the capillary to be supported only by the gel.

### 3D nucleus segmentation

3D nucleus segmentation of *Drosophila m*. embryos was performed using Ilastik [22]. The original dataset was compressed at different quantization levels, then upscaled in z to obtain isotropic resolution. To identify the nuclei, we used the pixel classification workflow, and trained it on the uncompressed dataset. This training was then used to segment the compressed datasets as well. Segmentation overlap was calculated in Matlab (Supplementary Code) using the Sørensen–Dice index [23], [24]:

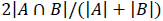

where the sets *A* and *B* represent the pixels included in two different segmentations.

### 3D membrane segmentation

Raw MuVi-SPIM recordings of *Phallusia mammillata* embryos expressing PH-citrine membrane marker were kindly provided by Ulla-Maj Fiuza (EMBL, Heidelberg). Each recording consisted of 4 views at 90 degree rotations. The views were fused using an image based registration algorithm followed by a sigmoidal blending of the 4 views. The fused stack was then segmented using the MARS algorithm [25] with an hmin parameter of 10. The raw data (all 4 views) was compressed at different levels, and segmented using the same pipeline. Segmentation results were then processed in Matlab to calculate the overlap score for the membranes using the Sørensen–Dice index (Supplementary Code).

### Single-molecule localization imaging

In order to visualize microtubules, U2OS cells were treated as in [26] and imaged in a dSTORM buffer [27]. In brief, the cells were permeabilized and fixed with glutaraldehyde, washed, then incubated with primary tubulin antibodies and finally stained with Alexa Fluor 647 coupled secondary antibodies. The images were recorded on a home-built microscope previously described [26], in its 2D single-channel mode.

### Single-molecule localization data analysis

Analysis of single-molecule localization data was performed on a custom-written MATLAB software as in [28]. Pixel values were converted to photon counts according to measured offset and calibrated gain of the camera (EMCCD iXon, Andor). The background was estimated with a wavelet filter [29], background-subtracted images were thresholded and local maxima were detected on the same images. 7-pixel ROIs around the detected local maxima were extracted from the raw images and fitted with a GPU based MLE fitter [30]. Drift correction was performed based on cross-correlation. Finally, images were reconstructed by filtering out localizations with a high uncertainty (>30 nm) and large PSF (>150 nm) and Gaussian rendering.

### Simulation of single-molecule localization data

Single molecule localization data was simulated in Matlab (Supplementary Code) by generating a grid of pixelated Gaussian spots with standard deviation of 1 pixel. With a pixel size of a 100 nm, this corresponds to a FWHM of 235.48 nm. The center of each spot was slightly offset from the pixel grid at 0.1 pixel increments in both x and y directions. To this ground truth image a constant value was added for illumination background, and finally Poisson noise was applied to the image. This process was repeated 10000 times to obtain enough images for adequate accuracy.

### Code availability

Source code and compiled binaries, including a filter plugin for HDF5, will be made available for download at https://git.embl.de/balazs/B3D. For immediate access, please send a request to.

## Acknowledgements

We thank Ulla-Maj Fiuza (European Molecular Biology Laboratory) for providing *Phallusia m*. datasets, Ulf Matti (European Molecular Biology Laboratory) for preparing samples for SMLM imaging, as well as Jan Ellenberg and Jean-Karim Hériché (both European Molecular Biology Laboratory) for providing screening microscopy datasets. The work was supported by the European Molecular Biology Laboratory, and the EMBL International PhD Programme (B.B, J.D. and M.A.). L.H. acknowledges support from the Center of Modeling and Simulation in the Biosciences (BIOMS) of the University of Heidelberg.

## Contributions

B.B. designed and implemented the compression algorithm and analyzed the data. L.H. and J.R. supervised the project. B.B. and M.A. performed light-sheet microscopy experiments. J.D. performed single molecule localization experiments and data analysis. B.B. wrote the manuscript with contributions from all other authors. All authors contributed to the final algorithm design.

## References

[1] A. E. Carpenter and D. M. Sabatini , “Systematic genome-wide screens of gene function,” Nat. Rev. Genet., vol. 5, no. 1, pp. 11–22, Jan. 2004.

[2] C. J. Echeverri and N. Perrimon , “High-throughput RNAi screening in cultured cells: a user’s guide,” Nat. Rev. Genet., vol. 7, no. 5, pp. 373–384, May 2006.

[3] R. Pepperkok and J. Ellenberg , “High-throughput fluorescence microscopy for systems biology,” Nat. Rev. Mol. Cell Biol., vol. 7, no. 9, pp. 690–696, Sep. 2006.

[4] J. Huisken , J. Swoger , F. Del Bene , J. Wittbrodt , and E. H. K. Stelzer , “Optical Sectioning Deep Inside Live Embryos by Selective Plane Illumination Microscopy,” Science, vol. 305, no. 5686, pp. 1007–1009, 2004.

[5] H.-U. Dodt et al., “Ultramicroscopy: three-dimensional visualization of neuronal networks in the whole mouse brain,” Nat. Methods, vol. 4, no. 4, pp. 331–336, Mar. 2007.

[6] U. Krzic , S. Gunther , T. E. Saunders , S. J. Streichan , and L. Hufnagel , “Multiview light-sheet microscope for rapid in toto imaging,” Nat. Methods, vol. 9, no. 7, pp. 730–733, Jul. 2012.

[7] R. Tomer , K. Khairy , F. Amat , and P. J. Keller , “Quantitative high-speed imaging of entire developing embryos with simultaneous multiview light-sheet microscopy,” Nat. Methods, vol. 9, no. 7, pp. 755–763, Jul. 2012.

[8] R. K. Chhetri , F. Amat , Y. Wan , B. Höckendorf , W. C. Lemon , and P. J. Keller , “Whole-animal functional and developmental imaging with isotropic spatial resolution,” Nat. Methods, vol. 12, no. 12, pp. 1171–1178, Dec. 2015.

[9] E. Betzig et al., “Imaging Intracellular Fluorescent Proteins at Nanometer Resolution,” Science, vol. 313, no. 5793, pp. 1642–1645, Sep. 2006.

[10] S. T. Hess , T. P. K. Girirajan , and M. D. Mason , “Ultra-High Resolution Imaging by Fluorescence Photoactivation Localization Microscopy,” Biophys. J., vol. 91, no. 11, pp. 4258–4272, Dec. 2006.

[11] M. J. Rust , M. Bates , and X. Zhuang , “Sub-diffraction-limit imaging by stochastic optical reconstruction microscopy (STORM),” Nat. Methods, vol. 3, no. 10, pp. 793–796, Oct. 2006.

[12] R. Wollman and N. Stuurman , “High throughput microscopy: from raw images to discoveries,” J. Cell Sci., vol. 120, no. 21, pp. 3715–3722, Nov. 2007.

[13] E. G. Reynaud , J. Peychl , J. Huisken , and P. Tomancak , “Guide to light-sheet microscopy for adventurous biologists,” Nat. Methods, vol. 12, no. 1, pp. 30–34, Jan. 2015.

[14] J. M. Perkel , “The struggle with image glut,” Nature, vol. 533, no. 7601, pp. 131–132, Apr. 2016.

[15] D. W. Cromey , “Digital Images Are Data: And Should Be Treated as Such,” Methods Mol. Biol. Clifton NJ, vol. 931, pp. 1–27, 2013.

[16] J. Nickolls , I. Buck , M. Garland , and K. Skadron , “Scalable Parallel Programming with CUDA,” Queue, vol. 6, no. 2, pp. 40–53, Mar. 2008.

[17] F. Amat , B. Höckendorf , Y. Wan , W. C. Lemon , K. McDole , and P. J. Keller , “Efficient processing and analysis of large-scale light-sheet microscopy data,” Nat. Protoc., vol. 10, no. 11, pp. 1679–1696, Nov. 2015.

[18] K. Sayood , Introduction to Data Compression, Fourth Edition, 4th ed. Morgan Kaufmann, 2012.

[19] T. Pietzsch , S. Saalfeld , S. Preibisch , and P. Tomancak , “BigDataViewer: visualization and processing for large image data sets,” Nat. Methods, vol. 12, no. 6, pp. 481–483, Jun. 2015.

[20] J. Schindelin et al., “Fiji: an open-source platform for biological-image analysis,” Nat. Methods, vol. 9, no. 7, pp. 676–682, Jul. 2012.

[21] G. de Medeiros et al., “Confocal multiview light-sheet microscopy,” Nat. Commun., vol. 6, p. 8881, Nov. 2015.

[22] C. Sommer , C. Straehle , U. Köthe , and F. A. Hamprecht , “Ilastik: Interactive learning and segmentation toolkit,” in 2011 IEEE International Symposium on Biomedical Imaging: From Nano to Macro, 2011, pp. 230–233.

[23] T. J. Sørensen , A method of establishing groups of equal amplitude in plant sociology based on similarity of species content and its application to analyses of the vegetation on Danish commons. København: I kommission hos E. Munksgaard, 1948.

[24] L. R. Dice , “Measures of the Amount of Ecologic Association Between Species,” Ecology, vol. 26, no. 3, pp. 297–302, Jul. 1945.

[25] R. Fernandez et al., “Imaging plant growth in 4D: robust tissue reconstruction and lineaging at cell resolution,” Nat. Methods, vol. 7, no. 7, pp. 547–553, Jul. 2010.

[26] J. Deschamps , M. Mund , and J. Ries , “3D superresolution microscopy by supercritical angle detection,” Opt. Express, vol. 22, no. 23, pp. 29081–29091, Nov. 2014.

[27] M. Heilemann et al., “Subdiffraction-Resolution Fluorescence Imaging with Conventional Fluorescent Probes,” Angew. Chem. Int. Ed., vol. 47, no. 33, pp. 6172–6176, 2008.

[28] J. Deschamps , A. Rowald , and J. Ries , “Efficient homogeneous illumination and optical sectioning for quantitative single-molecule localization microscopy,” Opt. Express, vol. 24, no. 24, pp. 28080–28090, Nov. 2016.

[29] I. Izeddin et al., “Wavelet analysis for single molecule localization microscopy,” Opt. Express, vol. 20, no. 3, pp. 2081–2095, Jan. 2012.

[30] C. S. Smith , N. Joseph , B. Rieger , and K. A. Lidke , “Fast, single-molecule localization that achieves theoretically minimum uncertainty,” Nat. Methods, vol. 7, no. 5, pp. 373–375, May 2010.

